# A homozygous genome-edited Sept2-EGFP fibroblast cell line

**DOI:** 10.1101/570622

**Authors:** Monika Banko, Iwona Mucha-Kruczynska, Christoph Weise, Florian Heyd, Helge Ewers

## Abstract

Septins are a conserved, essential family of GTPases that interact with actin, microtubules and membranes and form scaffolds and diffusion barriers in cells. Several of the 13 known mammalian septins assemble into nonpolar, multimeric complexes that can further polymerize into filamentous structures. While some GFP-coupled septins have been described, overexpression of GFP-tagged septins often leads to artifacts in localization and function. To overcome this ubiquitous problem, we have here generated a genome-edited rat fibroblast cell line expressing Septin 2 (Sept2) coupled to enhanced green fluorescent protein (EGFP) from both chromosomal loci.

We characterize these cells by genomic polymerase chain reaction (PCR) for genomic integration, by western blot and reverse transcriptase-PCR for expression, by immunofluorescence and immunoprecipitation for the colocalization of septins with one another and cellular structures and for complex formation of different septins. By live cell imaging, proliferation and migration assays we investigate proper function of septins in these cells.

We find that EGFP is incorporated into both chromosomal loci and only EGFP-coupled Sept2 is expressed in homozygous cells. We find that endogenous Sept2-EGFP exhibits expression levels, localization and incorporation into cellular septin complexes similar to the *wt* in these cells. The expression level of other septins is not perturbed and cell division and cell migration proceed normally. We expect our cell line to be a useful tool for the cell biology of septins, especially for quantitative biology.

## Introduction

The septins are a family of conserved cytoskeletal GTPases that perform essential functions in diverse processes such as cell division (Estey et al., 2010; Hartwell, 1971; Kinoshita et al., 1997; Spiliotis et al., 2005), ciliogenesis (Hu et al., 2010; Kim et al., 2010), tissue morphogenesis (Shindo and Wallingford, 2014) and the development of the nervous system (Ageta-Ishihara et al., 2013; Shinoda et al., 2010; Tada et al., 2007). They are involved in several forms of cancer (Angelis and Spiliotis, 2016; Calvo et al., 2015; Montagna et al., 2003; Russell et al., 2000; Verdier-Pinard et al., 2017), pathogen invasion (Dagdas et al., 2012; Mostowy et al., 2010; Nölke et al., 2016; Pfanzelter et al., 2018), neurodegenerative disease (Ihara et al., 2003; Kinoshita et al., 1998) and are the genetic basis for hereditary neuralgic amyotrophy (Bai et al., 2013; Kuhlenbäumer et al., 2005; Landsverk et al., 2009; Neubauer et al., 2018). The 13 mammalian septins from nonpolar, palindromic complexes assembled from several subunits into a rod-shaped monomer that can further polymerize into higher order structures that interact with membranes (Tanaka-Takiguchi et al., 2009; Zhang et al., 1999), actin (Kinoshita et al., 1997; Mavrakis et al., 2014; Smith et al., 2015) or microtubules (Nagata et al., 2003; Surka et al., 2002). While septins are considered the fourth cytoskeleton (Mostowy and Cossart, 2012), it’s many functions in biology are much less understood that those of most other cytoskeletal elements.

The creation of recombinant DNA constructs of genes tagged with the green fluorescent protein is an essential tool for the study of dynamic processes in cells. Since the beginning of work with recombinant septins, the overexpression of tagged septins has been reported to result in artifacts. The overexpression of Sept7-GFP leads to accumulation along axons on hippocampal neurons ((Xie et al., 2007) and our own observations) and the overexpression of Sept2-GFP/YFP leads to the formation of thick, curved filaments in MDCK cells (DeMay *et al*., 2011) and in NRK cells (Schmidt and Nichols, 2004a), raising doubts on the suitability of septin overexpression for the study of septin dynamics or interactions.

The development of genome-editing methods such as zinc-finger nucleases, TALENs and CRISPR/CAS has rendered the recombinant expression of gene constructs from the endogenous locus quickly accessible in mammalian cell systems. Indeed it was found for clathrin-mediated endocytosis that endogenously expressed proteins can differ functionally from overexpressed protein (Doyon et al., 2011). Since septins seem especially vulnerable to overexpression artifacts, we decided to generate a genome-edited cell line in which both alleles of the *Sept2* gene are endogenously tagged with the enhanced green fluorescent protein (EGFP) at the start codon. We thoroughly characterize the resulting homozygous clonal cell line for the expression of septins, the formation of complexes, colocalization of Sept2-EGFP with endogenous septins and cytoskeletal elements. We furthermore tested for defects in cytokinesis and cell migration and found no detectable differences between genome-edited and *wt* cells.

## Results

We aimed to use transcription activator-like effector nucleases (TALEN) for genome editing, as this enzyme-recognition based double-strand break generation leads to significantly less off-site effects than CRISPR/Cas9-based methods. To do so, we ordered NRK49F cells from the German collection of microorganisms and cell cultures (DSMZ), a near-diploid, spontaneously immortalized fibroblast cell line generated from rat embryo kidney tissue. We first sequenced the genomic *Sept2* locus (3q36) to find out, whether it agrees with the published rat genome sequence. We found that it does indeed overlap completely with the sequence available in the UCSC Genome Browser (http://genome.ucsc.edu/, gene number RGD: 620056) and moved on to find an optimal site for cutting with TALENs, which was found 7bp upstream the start codon of the *Sept2* gene. We designed TALENs for the induction of a double strand break at this position and used our genome-sequencing data to generate > 700 base pair long homology arms upstream of the *Sept2* gene and in the coding sequence of the *Sept2* gene around the site of the putative genomic double-strand break (Figure 1A, Figure S1). We cloned these homology arms around the coding sequence of *EGFP* to generate an integration matrix for insertion of *EGFP* into the coding sequence of Septin 2. We transfected the resulting plasmid together with plasmids encoding the left and right TALEN into NRK49F cells. The optimal temperature for TALE nucleases activity is 30 °C (Doyon et al., 2010) and we thus performed a cold shock treatment after transfection. After 10 days, we harvested cells and sorted them by FACS (fluorescence-activated cell sorting) to distinguish a population of cells expressing Sept2-EGFP from wild type (*wt*) cells. We found that about 0.2% of cells exhibited a detectable expression of EGFP (Figure S2). We collected this fraction and cultured the resulting cell population for one week before we FACS sorted them again into individual wells of 96-well plates. We found that now 98% of cells exhibited detectable EGFP expression, likely representing a population of heterozygous and homozygous cells with *EGFP* inserted into the *Sept2* locus on one or both chromosomes. To select for homozygous clones, we rationalized that these might be the brightest cells and thus sorted the top 6% of cells into 96-well plates at a density of 0.5 cells/well. We let the resulting clones grow for ten days and collected genomic DNA from them to analyze it by PCR. We found that most cells were heterozygous, exhibiting both bands for the *wt Sept2* gene and bands for the *Sept2* gene fused to the coding sequence of *EGFP*. However, we also found a few clones that exhibited only a single band for *Sept2-EGFP* (Figure 1B), suggesting that they were homozygous for the insertion. We then grew presumed homozygous and heterozygous clones and investigated them by western blotting for Sept2. We found that the clones that exhibited two bands for the genomic PCR also exhibited two bands in the western blot at positions consistent with the molecular size of Sept2 and Sept2-EGFP, demonstrating that they expressed both proteins. On the other hand, clones that exhibited but a single band for *Sept2-EGFP* in the genomic PCR exhibited a single band consistent with the size of Sept2-EGFP in the western blot as well (Figure 1C). We concluded that we indeed had generated both heterozygous and homozygous clones expressing Sept2-EGFP from the endogenous loci. Homozygous cells exhibited an even level of expression and Sept2-GFP localized similar to immunostaining of endogenous Sept2 in wt cells (Figure 1D). Sept2-EGFP localized to actin stress fibres but not to microtubules as expected for this cell type (Schmidt and Nichols, 2004a) (Figure 1E).

**Figure 1:**
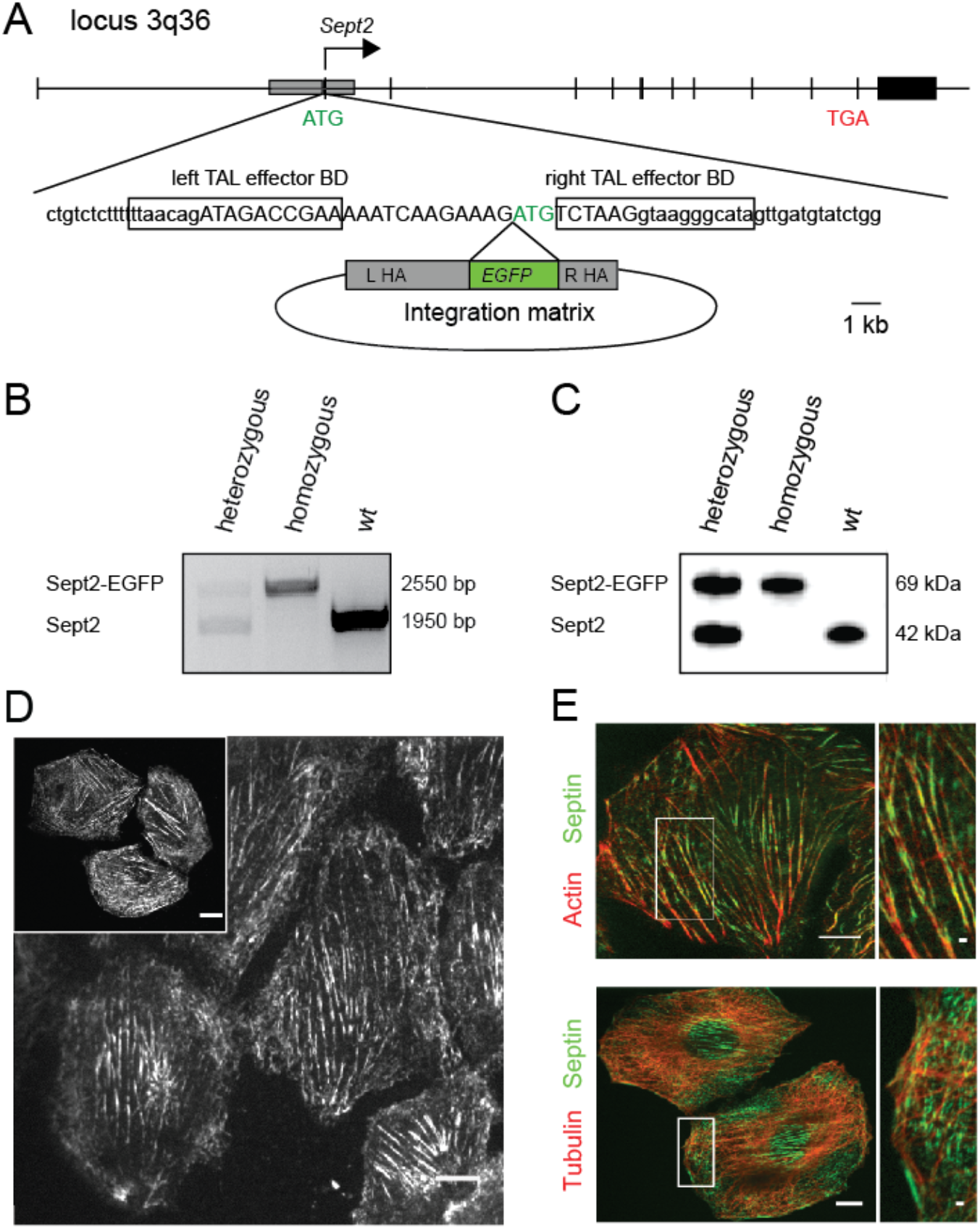
Design and initial characterization of a genome-edited NRK49F-Sept2 EGFP cell-line. **(A)** Strategy for the integration of *EGFP* into the rat *Sept2* locus. Exons shown in thick black. Recombination site used by the integration matrix represented by a grey box. Left and right TAL effector binding domains (BDs) enframed. *Sept2* exon given in the uppercase. The integration matrix contains left (LHA) and right (RHA) homology arms for homologous recombination. EGFP is inserted directly before and in frame with *Sept2* start codon (ATG, green). **(B)** Genomic PCR on the *Sept2* locus. Successful integration of *EGFP* into the *Sept2* locus results in a longer PCR product. Outcome for the wild type locus and single- and double-allelic integration shown. **(C)** Western Blot analysis on total cell extracts from wild type and genome-edited cells immunoblotted for Sept2. The same amount of protein was loaded into each lane. **(D)** Confocal microscopy image of live genome-edited NRK49F-Sept2-EGFP cells and fixed wild type NRK49F cells immunostained for Sept2 (inset). Scale bars are 10 µm. **(E)** Immunofluorescence micrograph of NRK49F-Sept2-EGFP cell-line showing EGFP fluorescence (green) and actin or tubulin staining (red). Sept2-EGFP decorates actin cables, but does not co-localize with tubulin in genome-edited NRK49F cells. Scale bars are 10 µm and 1 µm in the insets.

We next aimed to investigate the localization of endogenous Sept2-EGFP relative to other septins in the homozygous genome-edited cells. We first performed immunofluorescence staining against endogenous septins and performed colocalization analysis with Sept2-EGFP. We found that Sept7, Sept8, Sept9 and Sept11 colocalized with endogenous Sept2-EGFP (Figure 2A). When we then analyzed the homozygous cells by western blotting, we found that Sept2-EGFP, and, in agreement with the immunofluorescence data, septins 7,8,9,11 were detectable at the protein level (Figure 2B). Note that Sept2-GFP could be detected both via an anti-Sept2 as well as an anti-GFP-antibody. Sept6, however, could not be detected with all antibodies tested. When we tested *wt* cells, we detected the same septins at similar expression levels. Finally, we tested for expression by RT-PCR for all septins. While we could not detect Septins 3,4,12 and 14, we could detect all other septins, albeit at significantly different expression levels, with the septins detected by western blotting consistently being detected the strongest. Furthermore, the RNA expression of all endogenous septins was not significantly different between *wt* and genome-edited cells (Figure 2C). The only difference that may be detectable between *wt* and genome-edited cells is for Sept2-EGFP itself, which seems to be slightly higher expressed at the RNA-level in genome edited cells. The coding sequence for EGFP may stabilize the Sept2 mRNA. We concluded that septin expression was normal in our homozygous genome-edited Sept2-EGFP cell line.

**Figure 2:**
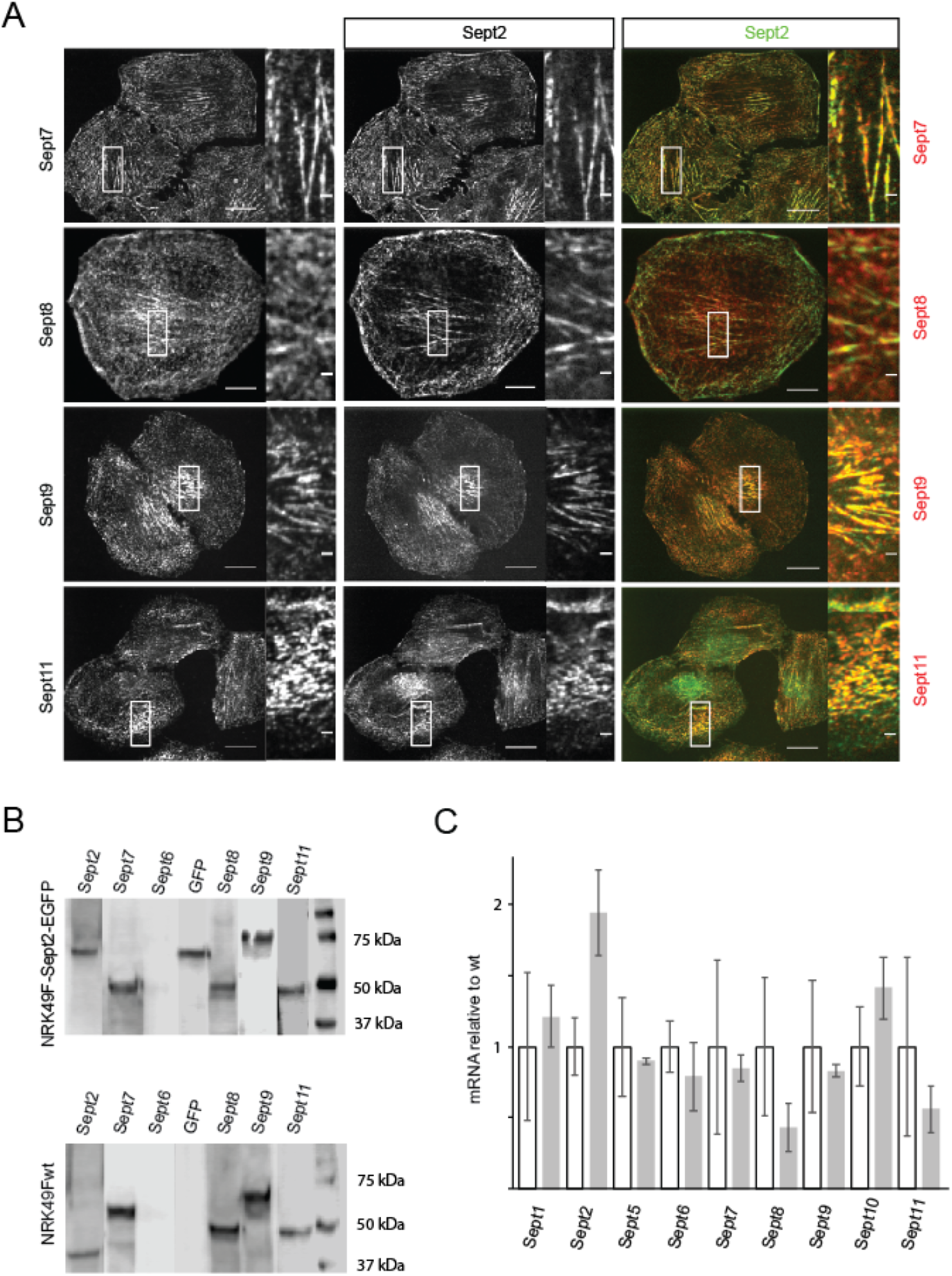
Septin expression in homozygous genome-edited NRK49F-Sept2-EGFP cells. **(A)** Immunofluorescence micrographs showing septin immunofluorescence staining (left), Sept2-EGFP fluorescence (right) and merged images. Scale bars: 10 µm and 1µm. **(B)** Western blot detection of septins in a total lysate from the genome-edited and wild type cell lines. **(C)** Real-time PCR analysis of cell lysate from *wt* and homozygous genome-edited cell lines. Shown is the detected mRNA level in homozygous genome-edited cell line relative to the *wt* expression level. Error bars are standard deviation.

We next aimed to determine, if Sept2-EGFP was incorporated into native complexes in genome-edited cells. To do so, we performed immunoprecipitation experiments using anti-GFP nanobody coupled to agarose beads. We found that Sept2-EGFP co-precipitated Sept7, Sept8, Sept9 and Sept11 (see Figure 3A). We achieved the same result when co-immunoprecipitating with anti-Sept2 antibodies, also in wt cells (data not shown). In contrast, when we performed immunoprecipitation experiments using GFP-nanobody beads in *wt* NRK49F cells, we could not detect any septins in the eluate, but only in full lysate (data not shown). When we performed mass spectrometry on septins immunoprecipitated via Sept2-EGFP purified form silver gels (Figure 3B), we could identify Sept7, Sept8, Sept9, Sept11. We concluded that Sept2 complex formation was not perturbed by the insertion of *EGFP* in the start codon in both alleles.

**Figure 3:**
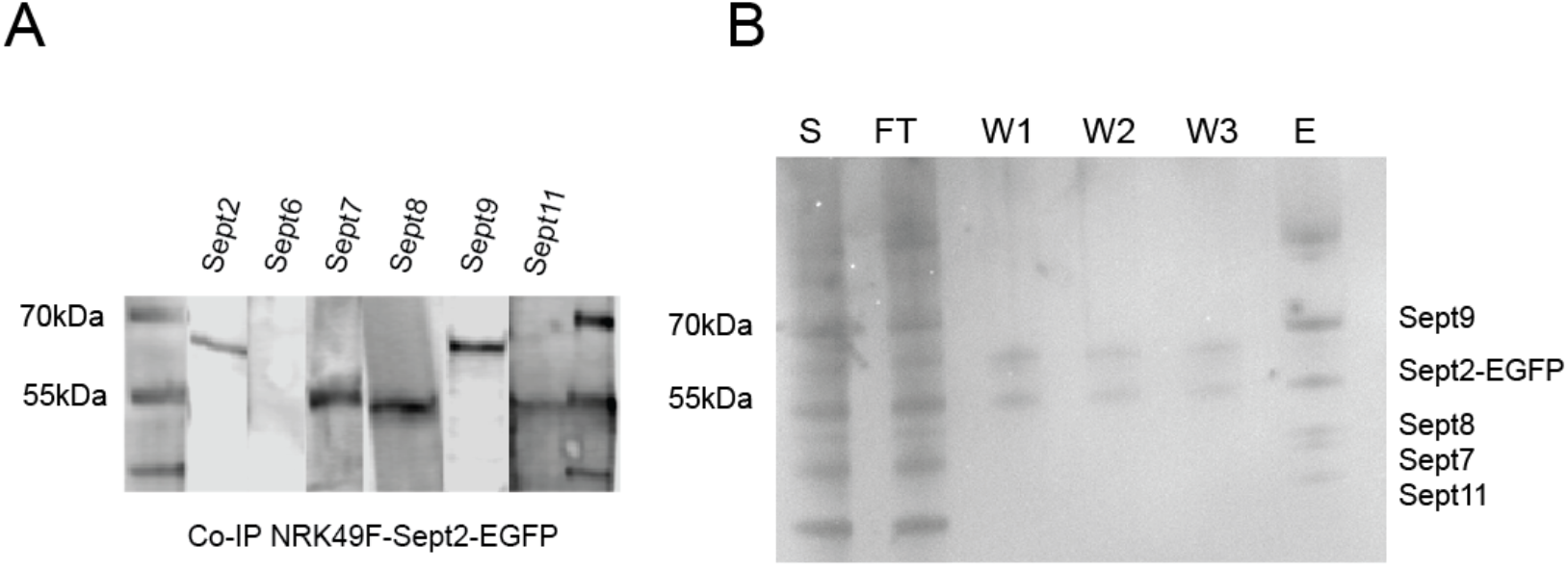
Co-immunoprecipitation of endogenous septins with Sept2-EGFP. **(A)** Western blot of septins co-immunoprecipitated from NRK49F-Sept2-EGFP cells using anti-GFP nanobody beads. **(B)** Silver-stained gel of immunoprecipitated septin complexes.

Since septins perform important functions in cell division, we next asked, whether septin localization was unperturbed during cell division. We performed immunofluorescence stainings of cells over the cell cycle in genome-edited and *wt* cells (Figure 4A) and found no difference in localization of septins over the cell cycle between *wt* and genome-edited cells. Similarly, when we observed dividing genome-edited cells in live-cell microscopy, we could observe normal distribution of Sept2-EGFP (Figure 4B). We concluded that Sept2 localization was not perturbed by the insertion of *EGFP* in the start codon in both alleles.

**Figure 4:**
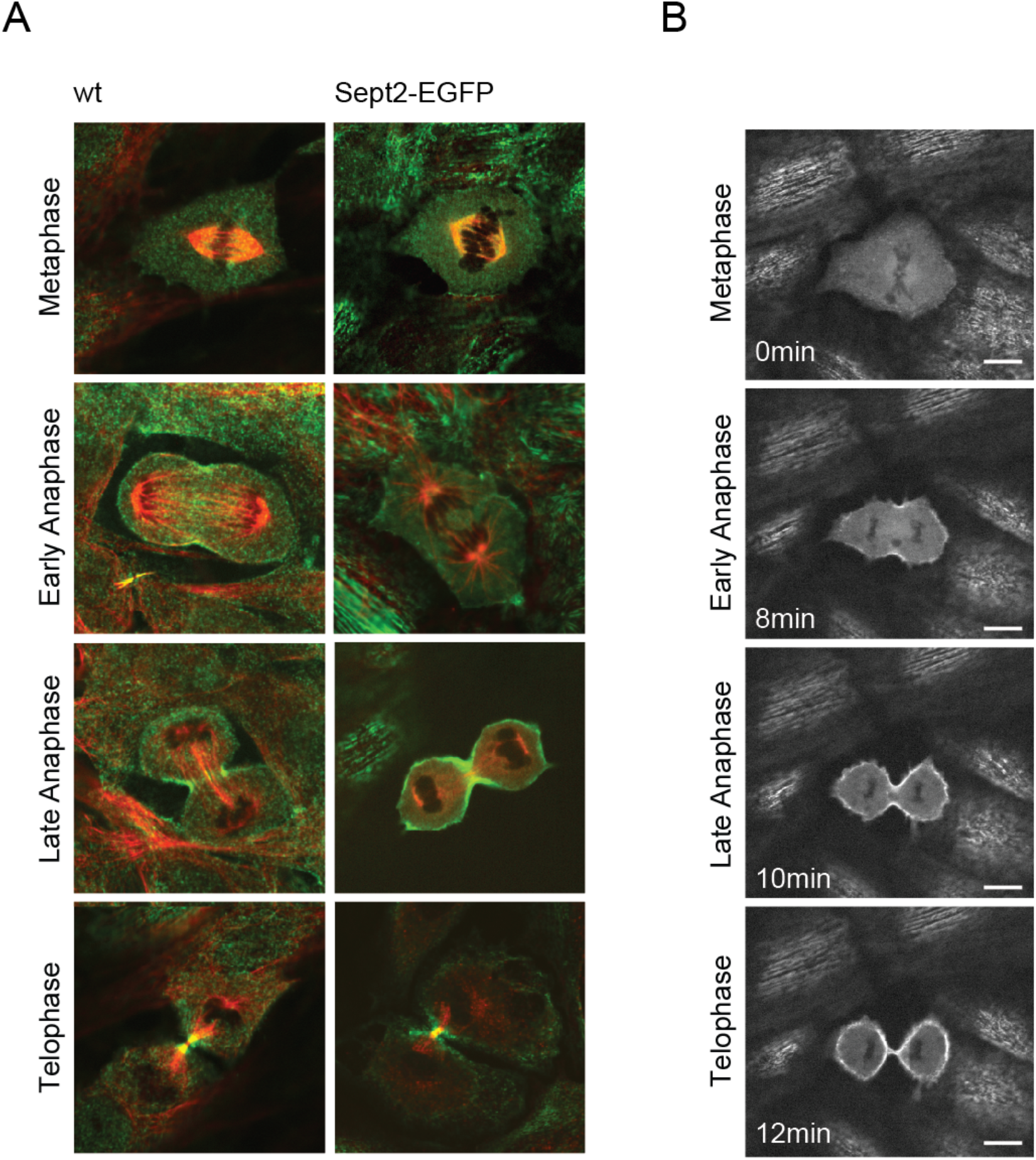
Comparison of Sept2 distribution during cell division in wild type and genome-edited cells. **(A)** Confocal images of wild type (left) and genome-edited (right) NRK49F fibroblast cells in different phases of the cell cycle. Sept2 antibody and Sept2-EGFP signal in green. Tubulin staining in red. **(B)** Individual frames from a live-cell time-lapse acquisition of dividing NRK49-Sept2-EGFP cells. Scale bars 10 µm.

Finally, we performed functional long-term assays for septin functions. We let cells divide for a number of cell cycles and analyzed for binucleated cells. Like in *wt* cells, we found in genome-edited cells only a minor fraction of around 2% of cells to be binucleated (Figure 5A). We also performed cell migration assays in form of wound healing assays. When we let cells migrate over 24h, we found no detectable difference in migration speed up to gap closure (Figure 5B). We concluded that cell division and cell migration are unperturbed in the genome-edited cells.

**Figure 5:**
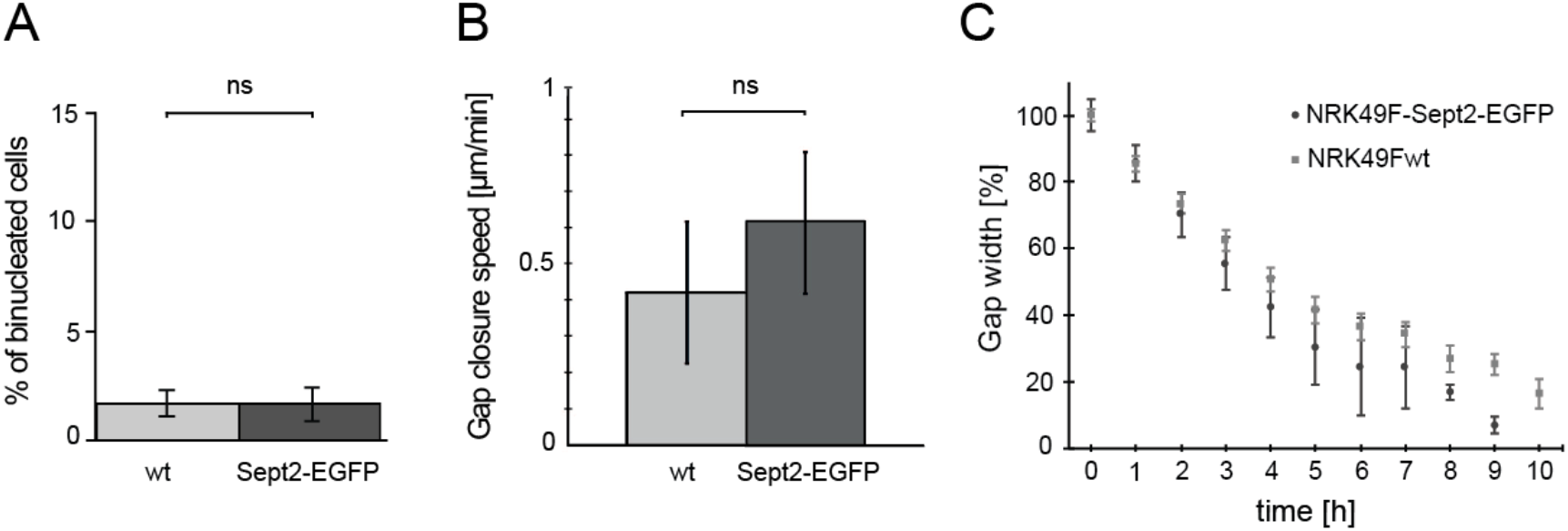
Cytokinesis and cell migration of the generated cells is not perturbed by the integration of *EGFP* into *Sept2* locus. **(A)** The average percentage of binucleated cells in wild type and genome-edited cells. **(B)** Quantification of migration speeds of wild type and genome-edited cells in a wound-healing assay. **(C)** Kinetics of wound closure for wild type and genome-edited cells in a wound-healing assay.

## Discussion

We have here generated a genome-edited cell line in which the coding sequence of the enhanced green fluorescent protein is incorporated at the start codon of both alleles of the *Sept2* gene. We have characterized this cell line thoroughly and found no detectable differences between *wt* cells and genome-edited cells. Since protein tagging with GFP has been established, artifacts have been reported for GFP-tagged molecules that were overexpressed via transient transfection of plasmids under strong promotors. Such artifacts may result from the titration of cellular interaction partners regulating complex formation, binding from subcellular localization or posttranslational modifications, or simple aggregation of the overexpressed molecule. The development of genome-editing techniques has lead to the possibility of tagging genes in mammalian cells at the genomic level in a straight-forward manner. Early studies have demonstrated a high potential of this technique to reduce artifacts, especially in dynamic processes (Doyon et al., 2011) and increasingly more cell biological studies rely on endogenous labeling. In the present work, we have generated a rat kidney fibroblast cell line that expresses Sept2-EGFP from the endogenous locus on both alleles. We chose this cell line, because rat is sequenced and has but one Sept2 splice isoform, NRK cells have been used in septin research before (Schmidt and Nichols, 2004b; 2004a) and they are diploid.

*Sept2* is an essential gene and its overexpression can lead to defects. This does not always need to be the case. It has been shown that overexpressed Sept7 can replace endogenous Sept7 that is downregulated via shRNA (Sellin et al., 2011), at least on the level of complex formation. In neuronal cells, shRNA-mediated downregulation of Sept5 can be functionally rescued by the overexpression of Sept2 and Sept4 from the same group (Kaplan et al., 2017), supporting the notion that exogenously expressed, tagged septins can be expressed and incorporated into functional complexes in cells. We find that Sept2-EGFP expressed from the endogenous locus does not lead to any detectable artifacts in localization, complex formation, or expression level of other septins. We furthermore find that cells that express but EGFP-tagged Sept2 divide and migrate similar to *wt* cells.

Taken together, we conclude that the cell line presented here will be a useful tool for septin biology and due to the homogenous, endogenous expression level will simplify the quantitative analysis of phenotypes observed in microscopy experiments. Especially in connection with single molecule localization based superresolution microscopy methods, this cell line will contribute to the advancement of our understanding of septin organization in cells (Kaplan et al., 2015; Ries et al., 2012; Vissa et al., 2018). More generally, it will simplify quantitative microscopy and microscopy-based screening and help with biochemical and proteomic approaches based on protein purification via the GFP tag (Trinkle-Mulcahy et al., 2008).

## Materials and Methods

### Cells

Rat kidney fibroblasts (NRK49F) were purchased from the German collection of microorganisms and cell cultures (DSMZ) and maintained in indicator-free Dulbeccos’s modification of Eagle’s medium (DMEM, Invitrogen) supplemented with 4.5 g/l Glucose, 100 mM Glutamax and 10% fetal bovine serum (Labforce). Cells were maintained in a humidified incubator with 5% CO_2_ at 37 °C.

### Genome-editing via TALENs

#### Genomic PCR

Genomic DNA was isolated with the GenElute mammalian genomic DNA miniprep kit (Sigma-Aldrich) according to the manufacturers protocol. The quality of isolated DNA was verified by agarose gel electrophoresis. The isolated DNA was used as a template to amplify the genomic sequence of Sept2 surrounding the START codon with the Sept2_genomicf and Sept2_genomicr primers, in a genomic PCR reaction under following conditions: denaturation 98 °C, 4 min; then 30 cycles of: 98 °C, 20 s; 61 °C, 20 s; 72 °C, 90 s and finally 10 min at 72 °C. The PCR products were analyzed by gel electrophoresis, and purified via the PureLink PCR purification kit (Invitrogen) according to the manufacturers protocol. The purified PCR products were sent to Microsynth AG for sequencing with primers Sept2_genomicf and Sept2_genomicr (Table 1).

**Table 1:**
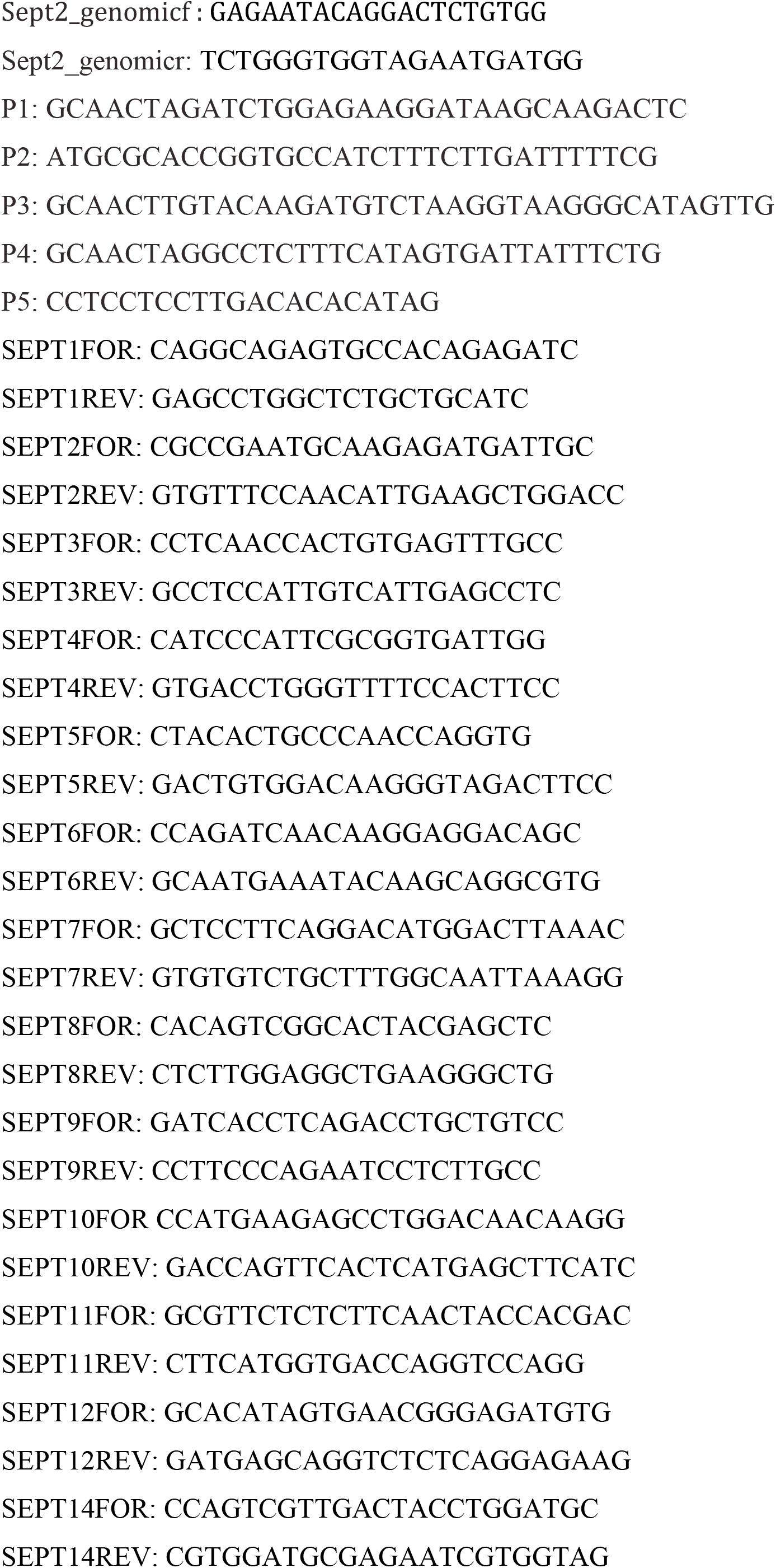
Primer sequences.

#### TALEN binding sequences

The pair of TALENs was designed and cloned by Cellectis bioresearch SAS according to the sequence of *Sept2* rat genomic DNA sequenced from NRK49F cells. The TALENs were design for a double strand break to occur 7 bp upstream of the start codon of the *Sept2* gene. The sequence recognized by the TALEN pair is underlined. The left and right TAL Effector DNA-binding domains are in bold print.

CACATACCTATCTTCATGGGTCAGTTCATGCTGTTAAATATGCATATACAA ATTCAGACTTCTGTCTCTCTTTT**<u>TTAACAGatagaccgaa</u>**<u>aaatcaagaaagATG**tctaagGTAAGGGCATA**</u>GTTGATGTATCTGGTGTAAAGAGAAATTATTTTAAATT TTCAGATCACTAGAAAAGATTTCTAGAGATAAATTTCTGATTTTGTATTCTAAAGTTTTGAGTTGCCATC

#### Generation of homology arms

The genomic DNA sequences surrounding the *Sept2* start codon, approximately 950 bp upstream from the START codon for the left homology arm (LHA) and 700 bp downstream for the right homology arm (RHA) were amplified using the P1/P2 and P3/P4 primers respectively (Table 1), genomic DNA isolated from NRK49F cells served as a template. The LHA was cloned into BglII and AgeI sites of the pEGFP-N1 vector (Clontech). The RHA was subcloned into BsrGI and StuI sites of the pEGFP-N1/LHA construct.

#### Transfection for genome-editing and clonal selection

Plasmids encoding for left and right TALENs and the homologous recombination donor plasmid (1.5 µg each) were transfected using a Neon Transfection system (Life Technologies), according to the manufacturer’s instructions, using 100 µl transfection tips. In brief, cells grown to ∼ 80 % confluency were harvested by trypsinization, washed twice in pre-warmed Ca2+/Mg2+-free PBS and 8 × 10^6^ cells were resuspended in 100 µl of room temperature solution R and electroporated with a pulse voltage of 1,650 V, a pulse width of 10 ms and pulse number of 3. After transfection, cells were incubated for 2 h at 37 °C and subjected to cold-shock treatment by incubation at 30 °C for 72 h before being transferred again to 37 °C, 5 % CO2. 5 to 7 days after the transfection, cells were FACS sorted for GFP-positive signals using a BD FACS AriaIIIu, 100 nozzle into a 6 cm dish and grown for 10 days. Amplified cells were re-sorted directly into 96-well plates (single cell per well) for clonal selection.

#### FACS sorting

Cells were harvested in 0.25 % Trypsin-EDTA. Trypsin was quenched with full DMEM (GIBCO) supplemented with 10% FCS and GlutaMAX (Invitrogen). Cells were then pelleted at 140 × g for 5 min. The supernatant was removed and cells were washed in Ca^2+^/Mg^2+^-free PBS (Invitrogen) and pelleted again at 140 × g for 5 min. The cellular pellet was gently resuspended in ice-cold, Ca^2+^/Mg^2+^-free PBS supplemented with 2 % FCS. Cells were then kept on ice and FACS sorted at 4 °C. The sorted cells were collected to pre-warmed full media and immediately transferred into 37 °C. Note, that decreasing the temperature during the sort from 37 °C to 4 °C and supplementing PBS with 0.2 % FCS significantly improved NRK49F and cell viability after the FACS sorting step.

#### Genomic DNA screening

Genomic PCR was used to screen for clones with the EGFP integrated on both alleles. Genomic DNA was isolated from expanded clones after the second FACS sorting step using the GenElute mammalian genomic DNA miniprep kit (Sigma-Aldrich) and amplified in a PCR reaction using primers P1/P5 (Table 1) under the following conditions: denaturation 98 °C, 4 min; then 30 cycles of: 98 °C, 20 s; 61 °C, 20 s; 72 °C, 90 s and finally 72 °C for 10 min. The size of PCR products were determined by gel electrophoresis to check for positive clones. The clones positive only for the modified *Sept2* gene were named NRK49F-Sept2-EGFP and their genomic DNA was verified by sequencing with primers P1 and P5.

### Preparation of cell lysate

For the verification of septin content in total cell lysates, 8 × 10^6^ cells were washed in ice cold PBS, lysed in BRB80 (80 mM PIPES pH 6.9, 1 mM EGTA, and 2 mM MgCl_2_) supplemented with complete EDTA-free protease inhibitor cocktail tablet (Roche) and 0.5 % saponin, harvested by scraping and homogenized by passing through a 25 Gauge needle. The cell lysates were mixed with 2 × concentrated sample buffer (126 mM Tris/HCl (pH 6.8), 20 % glycerol, 4 % SDS and 0.02 % bromophenol blue) and incubated at 95 °C for 7 min before loading on a SDS-PAGE gel.

### Co-immunoprecipitation and western immunoblotting

For co-immunoprecipitation we used GFP-trap magnetic beads (Chromotek) according to the manufacturers protocol. The cells were harvested in BRB80 (80 mM PIPES pH 6.9, 1 mM EGTA, and 2 mM MgCl_2_), complete EDTA-free protease inhibitor cocktail tablet (Roche)), supplemented with 0.5 % saponin. The lysis was performed on ice for 30 min; cells were then homogenized by passing through a 25 Gauge needle every 10 min. The homogenate was pre-clarified by centrifugation at 20’000 × g for 15 min at 4 °C. The post-nuclear supernatant was supplemented with NaCl to 0.5 M to prevent septin complex polymerization and spun at 200’000 × g for 40 min. Using this method, we have released above 90 % of all septins into the supernatant. Proteins were released from the beads with low pH according to the GFP-trap protocol, with 0.2 M glycine pH 2.5 and were neutralized with 1 M Tris base pH 10.4.

#### Antibodies used

Primary antibodies: Sept2 (SIGMA Atlas Antibodies HPA018481), GFP (Roche 12600500), Sept6 (Invitrogen PA5-42833, Sigma Atlas HPA003459, Santa Cruz Biotechnologies sc-20180, Proteintech 12805-I-AP), Sept7 (IBL JP18991), Sept8 (Santa Cruz Biotechnology sc-390105), Sept9 (SIGMA Atlas Antibodies HPA042564), Sept11 (Millipore ABN1342). All antibodies were used in concentration 1:1’000, except antibody against Sept7, which was diluted to 1:2’000. The blots were incubated overnight in 4°C with the antibodies on the shaking plate.

Secondary antibodies conjugated to horseradish peroxidase (HRP): Goat anti-rabbit (BioRad #170-6515), Donkey anti-mouse (Jackson Immunoresearch Laboratories 713-035-147). Secondary antibodies were diluted to 1:5’000 and incubated with blots for 30 min at room temperature on a shaking plate.

The proteins were detected with Amersham ECL Prime Western Blotting Detection Reagent (GE Healthcare, RPN2232), and visualized with LI-COR C-DiGit Chemiluminescence Western Blot Scanner.

### Immunofluorescence

For immunostaining, cells were fixed at RT in 4 % PFA, 2 % sucrose in BRB80 for 5 min. PFA was quenched in 50 mM NH_4_Cl/BRB80 for 10 min and washed 3 times with BRB80. Cells were permeabilized with 0.2 % TritonX-100 in 1 % BSA/ BRB80 for 4 min, and briefly washed in BRB80. For blocking we used 4 % horse serum in 1 % BSA/BRB80 for 45 min at RT. All septin antibodies were diluted to 1:1’000 in 1% BSA/BRB80 solution and incubated overnight at 4 °C. The primary antibodies were washed out 3 times for 5 min each on a 160 rpm shaking plate. To visualize septins, we used either Alexa647, Alexa488 or Alexa565 goat anti-rabbit secondary antibodies (from Invitrogen) diluted 1:800 in 1 % BSA/BRB80 and incubated them with cells for 30 min at RT. The secondary antibodies were washed out 3 times for 5 min each on a 160 rpm shaking plate.

#### Primary antibodies used for immunostaining

Sept2 (SIGMA Atlas Antibodies HPA018481), Sept7 (IBL JP18991), Sept8 (kind gift from Koh-Ichi Nagata), Sept9 (SIGMA Atlas Antibodies HPA042564), Sept11 (kind gift from Bill Trimble), Monoclonal-Anti-β-Tubulin-Cy3 (SIGMA) (1:5’000). To visualize actin we used Phalloidin ATTO-565 (1:10’000).

### Co-immunoprecipitation and mass spectrometry of septin complexes

NRK49F-Sept2-EGFP cells were lysed in BRB80 buffer (80 mM PIPES, 4 mM EGTA, 2 mM MgCl_2_, pH 6.9) supplemented with 0.5 % saponin and an EDTA-free protease inhibitor cocktail (Roche). Lysates were kept on ice for 30 min and drawn 15 times through 25 Gauge syringe needle every 10 min. Obtained homogenate was pre-clarified by centrifugation at 20 000 × g for 25 min. The supernatant was supplemented with NaCl to 0.5 M and incubated with equilibrated GFP-Trap magnetic beads (Chromotek).

After 15 h incubation at 4 °C, beads were washed 3 times with BRB80-0.5 M NaCl buffer, resuspended in 2 × SDS sample buffer, boiled at 95 °C for 10 min and magnetically removed. Isolated proteins were separated by sodium dodecyl sulfate-polyacrylamide gel electrophoresis (SDS-PAGE) and stained with Coomassie R-250. The gel bands were excised, reduced, alkylated, and digested with trypsin (Pierce 90058) in 25 mM ammonium bicarbonate overnight at 37 °C as described previously (Shevchenko et al., 2006). Samples were spotted using the dried-droplet technique onto a-cyano-4-hydroxycinnamic acid Maldi matrix. Spectra were obtained using Bruker Ultraflex II Maldi TOF/TOF Mass Spectrometer (Bruker Daltonics, Bremen, Germany) equipped with a 200 Hz solid-state Smart beam laser. The MS/MS spectra of selected peptides were acquired in the LIFT mode. Mascot data base search with the peptide mass fingerprint data obtained from the top band identified a mixture of septins 2 and 9, and from the lower band a mixture of septins 7, 8, and 11. Since the score for septin 8 was only slightly above the significance threshold, this identification had to be confirmed by two MS/MS spectra that yielded unique sequences for septin 8 (m/z=1021.5, EFLSELQR = SEPT8_RAT pos. 332-339; m/z=1941.0, LRPQTYDLQESNVHLK = SEPT_RAT pos. 083-098) proving unambiguously its presence in the septin complex.

### RT-PCR

RNA from NRK49F-*wt* and NRK49F-Sept2-EGFP cells were extracted using ‘RNA Tri flüssig’ (Bio & Sell) according to the manufacturer’s protocol. RNA was reconstituted in ddH2O and adjusted to 0.5 µg/µl. For RT-PCRs, 1 µg RNA per reaction were reverse transcribed using random hexamers and M-MuLV reverse transcriptase in 20 µl reactions using standard protocols. cDNA was then diluted to 200 µl total volume and processed in qPCR. qPCR was performed using the PowerUp SYBR Green Master Mix (Thermo Fisher Scientific, A25742) in a 20 µl reaction according to the manufacturer’s protocol.

### Determination of number of binucleated cells

Genome-edited or *wt* cells (1 × 10^5^) were seeded on 18 mm coverslips. 48 h after seeding, the cells were fixed in 4 % PFA, 2 % sucrose in BRB80, permeabilized in 0.2 % Triton X-100 in 1% BSA/BRB80 and stained with Phalloidin-ATTO488. To visualize the nucleus, the coverslips where mounted in Vectashield H-1000 (Vector Laboratories, Inc. CA, USA) mounting medium for fluorescence containing propidium iodide. The number of cells with single and well-separated double nuclei was counted (n > 400 cells) during three independent experiments in triplicate.

### Migration assay

A confluent monolayer of wild type and genome-edited cells seeded on poly-l-lysine coated coverslips was linearly wounded with a 10 µl pipette tip. Cells were gently washed three times in BRB80 or PBS and either fixed in 4 %PFA/BRB80 as described above or imaged in DMEM full media supplemented with Pen/Strep antibiotic for long-term live-cell imaging. Time-lapse movies of migrating cells were recorded in phase contrast on a Biostation IM-Q (Nikon) using a 10x objective. Images of fixed positions in the wounds were taken with intervals of 10 min for 10 h. Through the experiment, cells were incubated in full DMEM supplemented with 2 % FCS, 100 U/ml penicillin and 100 U/ml streptomycin (Invitrogen). Using NIH ImageJ analysis software, and a reference picture of a micrometer slide, the distances between the edges of the wound were recorded over time. The average distance between the wound edges in µm were plotted against time in Microsoft Office Excel software, and a linear fit was generated for each dataset. The slope of the linear fit was used as a measure of cell migration speed.

### Live-Cell imaging

Cells were kept in indicator-free Dulbeccos’s modification of Eagle’s medium (DMEM, Invitrogen) supplemented with 4.5 g/l Glucose, 100 mM Glutamax and 10% fetal bovine serum (Labforce) and 50 mM HEPES, pH 7.0. Cells were seeded on 18 mm coverslips at density 1 × 10^5^ and imaged on a spinning disc confocal microscope equipped with a 100 × Olympus (PlanSApo N, NA 1.40 NA oil) objective. The sample holder stage was heated to a temperature of 37 °C and images were taken in 10 min intervals.

## Acknowledgments

This work was supported by a Muller Fellowship (M.B.), ETH Zurich, King’s College London, FU Berlin, the Deutsche Forschungsgemeinschaft (DFG, German Research Foundation) - Project Number 278001972 - TRR 186 and SFB 958. We thank Bill Trimble and Koh-Ichi Nagata for antibodies and all present and former members of the Ewers laboratory for helpful discussions.

## Supplementary data

**Figure S1:**
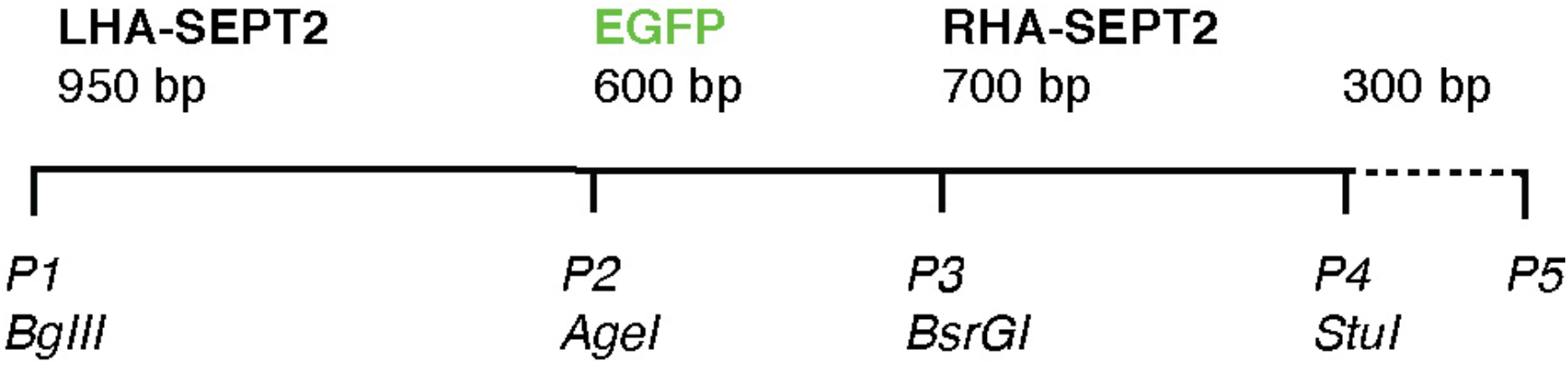
Matrix used for EGFP insertion into the *Sept2* locus by homologous recombination. Chromosomal DNA outside of the integration matrix is indicated by a dashed line. The left homology arm (LHA-Sept2) is a 950 bp fragment of genomic DNA upstream of the *Sept2* start codon and the right homology arm (RHA-Sept2) is a 700 bp fragment downstream from the start codon. Both fragments are complementary to the chromosomal *Sept2* sequences around the double strand break. Integrated fluorescent tag fragment marked in green. The primer sites *P1-P7* with the indicated restriction sites were used for cloning and sequencing. *P5* binds downstream of the sequence contained in the integration matrix and was used together with *P1* for screening for clones with a successfully integrated EGFP.

**Figure S2:**
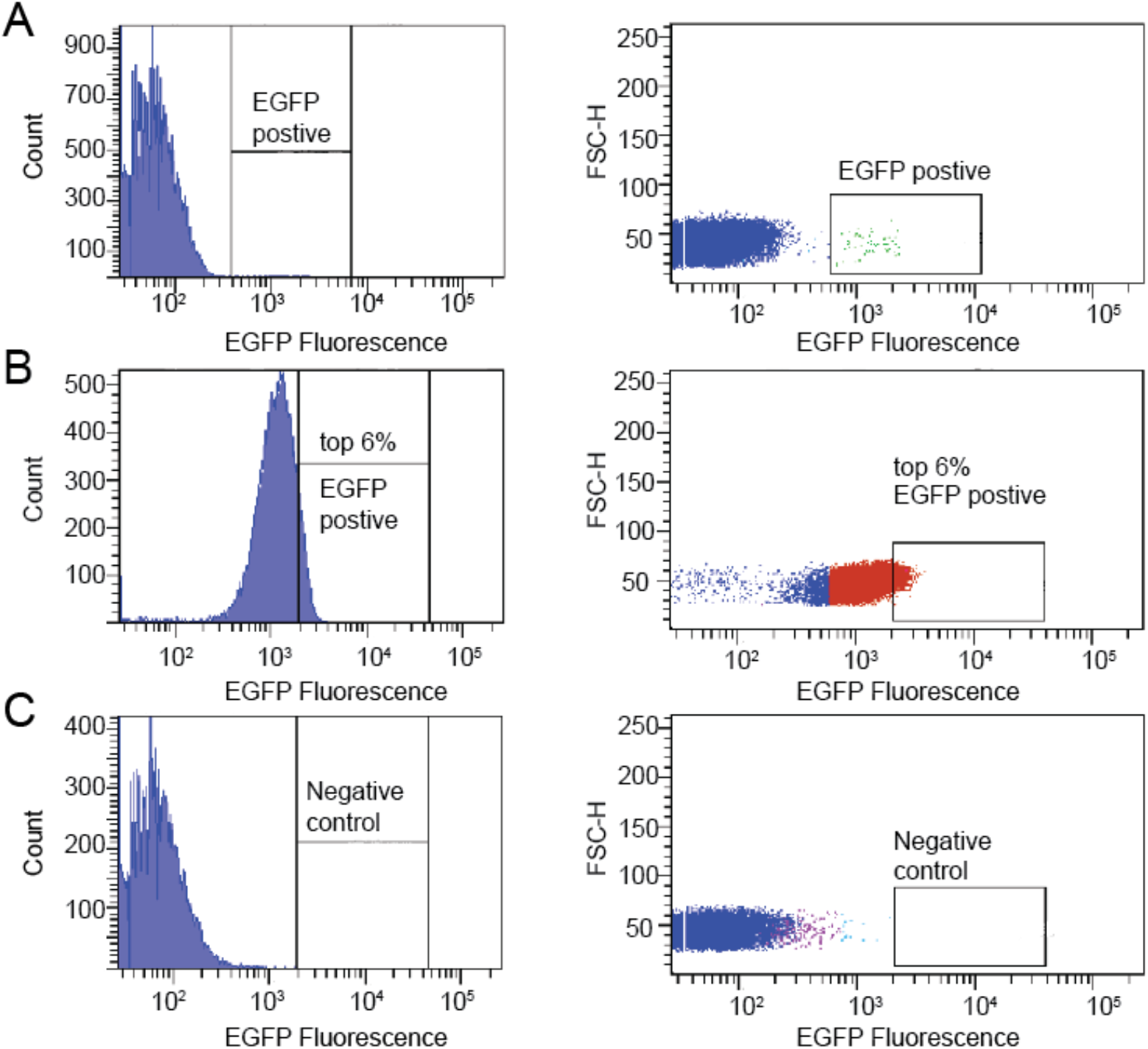
Dot plots and histograms of FACS sorted NRK49F cells transfected with TAL effector nucleases and the EGFP integration matrix. **(A-C)** Histograms (left) and dot plots (right) show EGFP fluorescence distribution. Sorting gates for EGFP positive cells are marked in boxes. **(A)** First FACS sorting of NRK49F cells transfected with TAL effector nucleases and an integration matrix with EGFP. **(B)** Second FACS sorting of single cells expressing EGFP. Note, that for a single cell sorting only the cells in the top 6% of the EGFP fluorescence were collected. **(C)** NRK49F cells not transfected with the integration matrix served as a negative control.

